# *Withania somnifera* showed neuroprotective effect and increase longevity in *Drosophila* Alzheimer’s disease model

**DOI:** 10.1101/2020.04.27.063107

**Authors:** Mardani Abdul Halim, Izzah Madihah Rosli, Siti Shafika Muhamad Jaafar, Hoi-Min Ooi, Pui-Wei Leong, Shaharum Shamsuddin, Nazalan Najimudin, Ghows Azzam

## Abstract

Alzheimer’s disease is a complex neurodegenerative disease and is only unique to human. The disease is defined in human brain by the accumulation of amyloid beta in the parenchyma of the brain. Withania somnifera, commonly known as Ashwagandha is an Indian Ayurvedic medicine that has been used for centuries to treat countless range of human health problem. The active compound of Ashwagandha was shown to be beneficial in treating many neurodegenerative diseases including Alzheimer’s disease (AD). In this study, *Drosophila* melanogaster AD model was used to study the effect of Ashwagandha on the toxicity of beta amyloid and also the longevity effect of the compound. We found that 20 mg/mL of Ashwagandha was shown to be effective in rescuing the “rough eye phenotype” of AD *Drosophila*. Furthermore, Ashwagandha also promotes longevity in AD as well as wild-type *Drosophila*. The results above showed that Ashwagandha could potentially be a potent drug to treat AD as well as maintaining the wellbeing of cells.

## 1. Introduction

Alzheimer’s disease (AD) can be described as progressive neurodegenerative disorder that affect wide range of the cerebral cortex and hippocampus. A number of hypotheses have been suggested in order to explain the pathogenicity of AD like oxidative stress theory, amyloid hypothesis and also APOE genotype hypothesis (1–3). The primary cause of AD is the accumulation of amyloid-β (Aβ) peptide in which the will contributed to the oxidative injury as well as inflammation to the brain (4).

There are two classes of drugs that have been approved to treat AD namely donepezil, tacrine, galantamine and rivastigmine in which are acetylcholinesterase inhibitors. This class of drug protects neurotransmitter acetylcholine against breaking down. Another class is the N-methyl-d-aspartate receptor, memantine in which it blocks the action of neurotransmitter glutamate where it can cause neuron death. However, both drugs incapable of stopping the progression of neuritic dystrophy (4) and thus alternative drug to treat AD is essential.

*Withania somnifera* (Ashwagandha) also known as Indian ginseng, is a sub-tropical herb originated in India has been recognized to boost neuronal function, reduce stress and anxiety as well as promote cerebral calmness (5). Major therapeutic component of Ashwagandha is withanolides, naturally occurred C-28 steroidal lactones. Countless studies have recognised Ashwagandha as an alternative strategy in treating several neurological disorders including Parkinson’s disease, Huntington’s disease, cerebral ischemia and also Alzheimer’s disease (6). In this study, we use *Drosophila melanogaster* as model to study the effect of Ashwagandha against lifespan of Alzheimer’s disease flies.

## 2. Materials and methods

### 2.1 Fly rearing and husbandry

All D. melanogaster stocks used in this study are listed in Flybase (http://flybase.bio.indiana.edu). Oregon-R wild-type (genotype: Oregon-R-P2; stock no.: 107294), Act5C-GAL4 (genotype: y[1] w[*]; P{w[+mC]=Act5C-GAL4}25FO1 / CyO, y[+]; stock no.: 107727) and GMR-GAL4 (genotype: w[*];P{w[+mC]=GAL4-ninaE.GMR}12; stock no.: 106207) were from Kyoto Stock Center and UAS-Aβ42 (genotype: w[1118];P{w[+mC]=UAS-APP.Abeta42.B}m26a; stock no.: 33769) was from Bloomington *Drosophila* Stock Center. All stocks were maintained at 25 °C in a corn-based meal consists of 4% (w/v) corn starch, 5% (w/v) polenta, 10% (w/v) brown sugar, 0.7% (w/v) agar, 5% (w/v) yeast, 3% (w/v) nipagin and 0.7% (v/v) propionic acid. All targeted expression was performed using Act5C-GAL4 and GMR-GAL4. For control, Oregon-R was crossed with GMR-GAL4 and Act5C-GAL4 to produce GMR-OreR and Act5C-OreR. For flies that expressed Aβ42 gene, GMR-GAL4 and Act5C-GAL4 was crossed with UAS-Aβ42 to produce transgenic GMR-Aβ42 and Act5C-Aβ42 respectively.

### 2.2 Treating AD flies with Ashwagandha

To generate AD flies, male of GMR-GAL4 (parent) was crossed with virgin female of UAS-Aβ42 (parent) and was kept in corn-based meal supplemented with 20 mg/mL Ashwagandha (root extract) (Himalaya, India). Seven days post-mating, the parents were removed and the pupae were left until F1 emerged. The F1 will express the Aβ42 protein in their compound eyes leading to rough eye phenotype (REP). A total of three types of crosses were conducted, GMR-GAL4 x UAS-Aβ42 (treated with Ashwagandha), GMR-GAL4 x UAS-Aβ42 (untreated) and GMR-GAL4 x OreR (control). All experiments were done in triplicates. The degree of REP was evaluated for all F1 to determine the effect of Ashwagandha.

### 2.3 Longevity assay

Longevity assay was performed to observe the effect of Ashwagandha towards the longevity of AD flies. In this experiment, UAS-Aβ42 was crossed with ubiquitous driver Act5C-GAL4. Parents were mated in corn-based meal supplemented with 20 mg/mL Ashwagandha. The emerged F1 was also being kept in the same food for the whole experiment. The total number of F1 emerged for each day was recorded until 100 F1’s were collected. The F1 was transferred into new food supplemented with compound in every three days. The number of death was recorded daily until all the F1 was dead. The data obtained was converted into a longevity graph. A total of three types of crosses were conducted, Act5C-GAL4 x UAS-Aβ42 (treated with Ashwagandha), Act5C-GAL4 x UAS-Aβ42 (untreated) and Act5C-GAL4 x OreR (control). All experiments were done in triplicates. In addition, OreR was also subjected to longevity assay in order to see the effect of Ashwagandha on wild-type flies.

## 3. Results

The compound eye of *Drosophila* composed of approximately 800 repeating subunits called ommatidia. Each ommatidium equipped with eight photoreceptor neurons. Ectopic expression of human Aβ42 under GMR-GAL4 driver (GMR-Aβ42) in *Drosophila* compound eyes resulted in a vibrant neurodegenerative phenotype in which projected by the disruption of the neurons (Figure 1A). The ommatidium orientation was distorted and thus giving the “rough eye phenotype”. On the other hand, GMR-Aβ42 Ashwagandha-treated flies showed significant improvement on their compound eyes compared to the untreated flies (Figure 1B). The ommatidium showed less “rough eye phenotype” as well as more organized compared to the untreated flies. The results showed that Ashwagandha has the ability to treat neurodegenerative disease cause by Aβ42 toxicity. For control, the ommatidium was perfect in shape and organization (Figure 1C).

**Figure 1.**
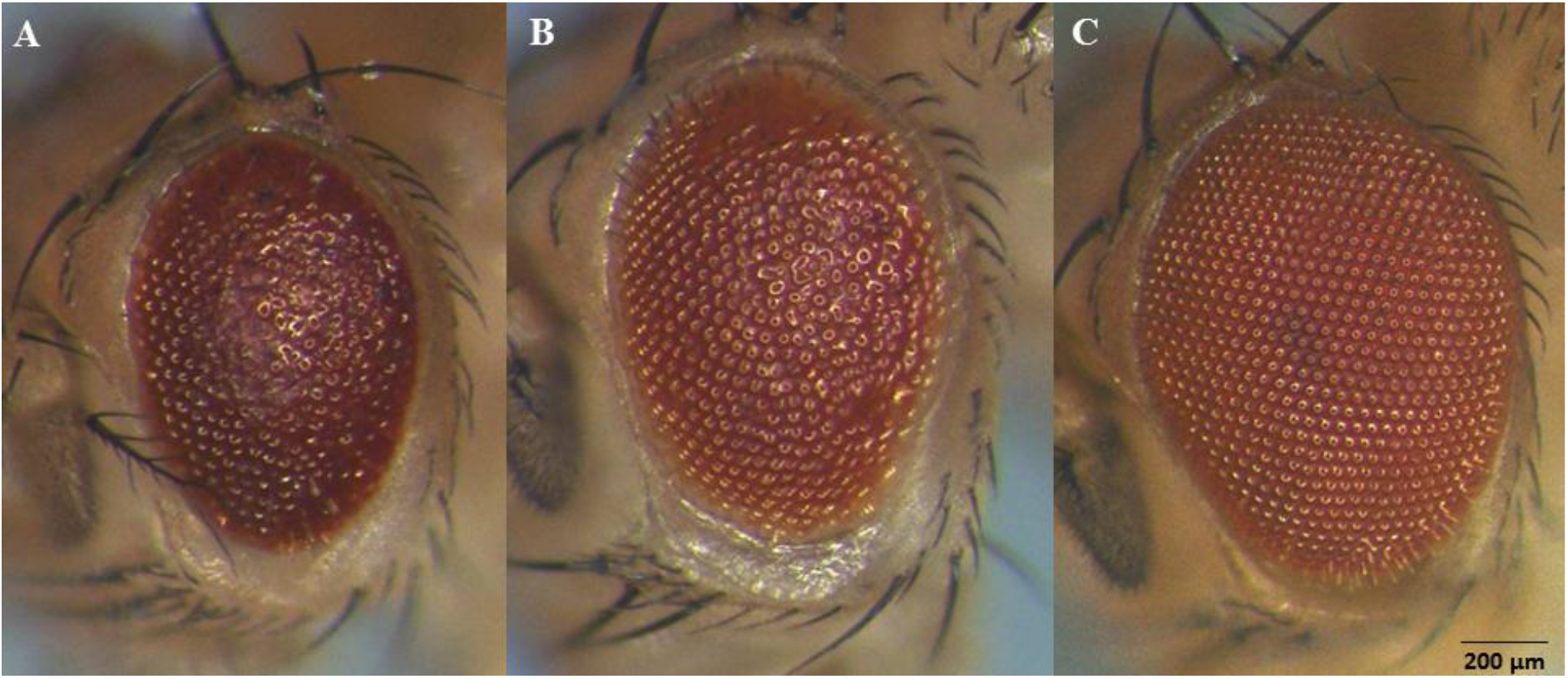
Rough eye phenotype of *Drosophila.* **A:** GMR-Aβ42 untreated flies. The ommatidia was irregular in shape. Fused omatidium was also observed as a result from amyloid beta toxicity. **B:** GMR-Aβ42 Ashwagandha-treated flies. The ommatidia was significantly improved compared to A. Less fused-ommatidium was observed due to the treatment of Ashwagandha. **C:** GMR-OreR (control). The ommatidia was perfectly organized with no fused-ommatidium observed.

To see if Ashwagandha could rescue the effect of premature death in AD flies, we tested the compounds using the longevity assay. For the longevity assay, Act5C ubiquitous driver was used. The untreated Act5C-Aβ42 flies live for approximately 16 days, while the Act5C-Aβ42 treated with Ashwagandha showed improvement in their lifespan living up to 29 days which is almost doubled from the untreated flies (Figure 2A). Interestingly, the same phenomenon was also observed on OreR treated with Ashwagandha. The treated flies showed increment in their lifespan in which the lived 5 days longer compared to the untreated OreR flies (Figure 2B). The results demonstrated that Ashwagandha has the property to promote longevity to both AD and wild-type flies.

**Figure 2.**
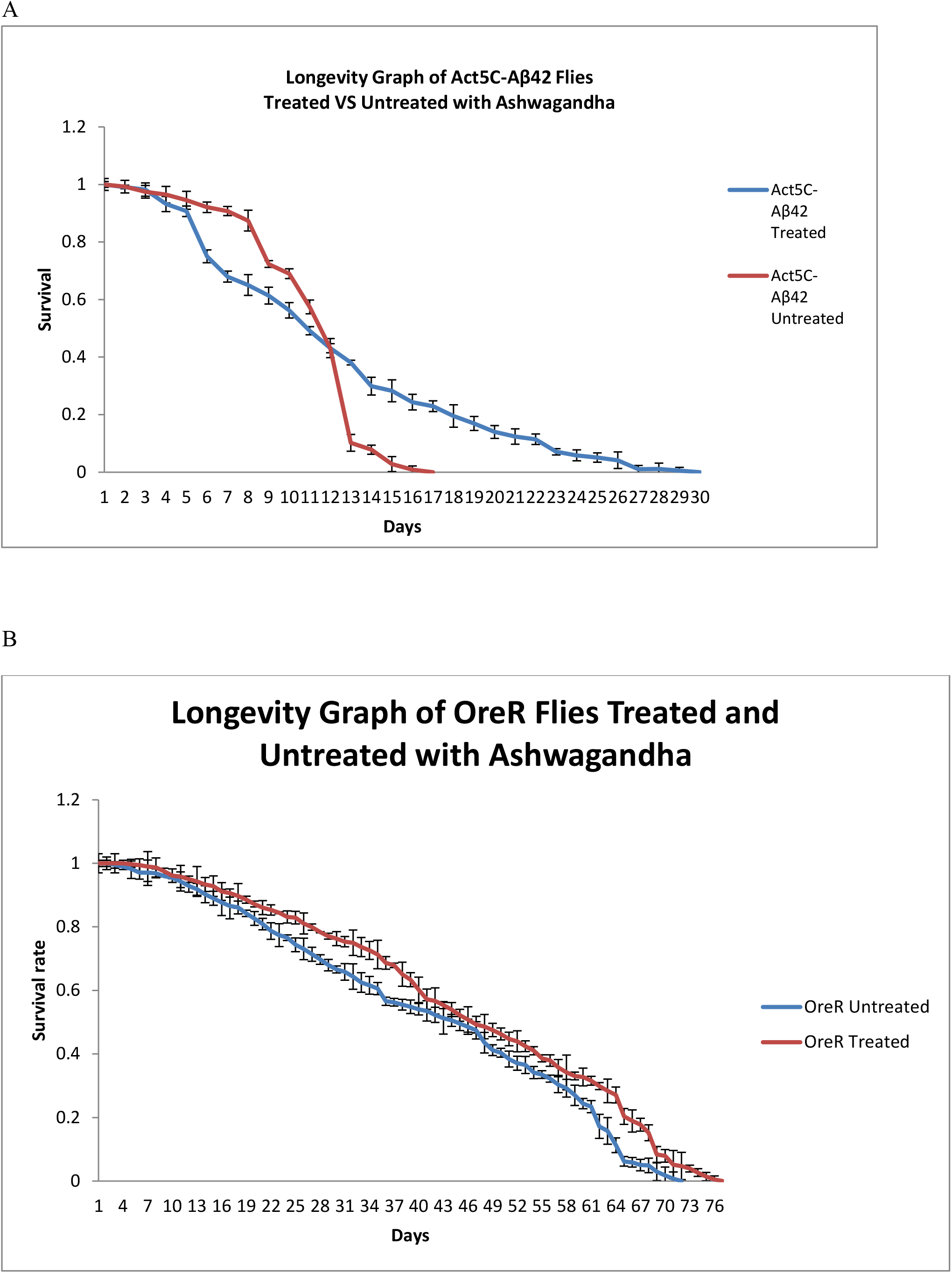
Longevity graph of *Drosophila*. **A:** Longevity graph of Act5C-Aβ42 flies treated versus untreated with Ashwagandha. The treated flies live significantly longer compared to the untreated flies. The untreated flies live approximately 16 days while Ashwagandha-treated flies live around 29 days. **B:** Longevity graph of wild-type OreR flies treated versus untreated with Ashwagandha. The treated flies live around 76 days while the untreated flies live around 71 days.

## 4. Discussion

Withanolides are the active compounds in Ashwagandha that are responsible for countless biological activities. AD can be characterized by extracellular deposit of amyloid beta protein as well as intracellular neurofibrillary tangles in the cortical and limbic regions of the brain. The use of *Drosphila* REP as an indicator of neuronal damage caused by amyloid beta protein is the simplest way to screen for potential drugs to combat AD. Previous study showed that Ashwagandha can induces the reversal of amyloid beta toxicity in SK-N-MC cells and also increased acetylcholinesterase activity in neuronal cell culture (7). Active compound in Ashwagandha was also shown to down-regulate beta-secretase 1 (BACE1) and up-regulating disintegrin and metalloprotease 10 (ADAM10) in amyloid beta clearance (8). In this study, the clearance of amyloid beta in *Drosophila* is mediated probably by the two mechanisms aforementioned. Another possible mechanism was the reversal of beta amyloid accumulation peptides and oligomers as shown by Seghal et al. in 2012 (9).

Longevity assay was performed in order to see the effect of Ashwagandha on the lifespan of AD as well as wild-type flies. From the result, AD flies treated with Ashwagandha showed significant increase in their lifespan compared to the untreated AD flies. Although the mechanism of Ashwagandha mediated longevity in *Drosophila* still unknown, there are several other studies on other organisms that might have similar mode of action. For instance, Ashwagandha was found to extend approximately 20% of the lifespan of human EGFR-driven cancerous *Caenorhabditis elegans* by targeting the Insulin/IGF-1 signalling pathway (10). Besides, Ashwaganda treatment also showed around 20% lifespan extension in human nicotinic receptor, nAchR*, α7* equivalent, *acr-16* mutant *C. elegans.* Surprisingly, there was no effect observed on wild type *C. elegans* (11). This was totally opposite from our data where wild-type *Drosophila* also showed lifespan extension around 5% when treated with Ashwagandha compared to the untreated flies.

## Conclusion

This study reveals that Ashwagandha treated AD *Drosophila* showed significant improvement in their REP compared to the untreated flies. Moreover, Ashwagandha also promotes longevity in AD as well as wild-type flies. However, the mechanisms behind the longevity effect on wild-type flies still remain unknown.

## Acknowledgement

We would like to thank all our collaborators and colleagues for the discussion and the work conducted in this lab. This study was funded by the USM Top Down Research Fund – URICAS (1001/PBIOLOGI/870029).

